# Heartbeats entrain breathing via baroreceptor-mediated modulation of expiratory activity

**DOI:** 10.1101/2020.12.09.416776

**Authors:** William H. Barnett, David M. Baekey, Julian F. R. Paton, Thomas E. Dick, Erica A. Wehrwein, Yaroslav I. Molkov

## Abstract

Cardio-ventilatory coupling refers to a heartbeat (HB) occurring at a preferred latency before the onset of the next breath. We hypothesized that the pressure pulse generated by a HB activates baroreceptors that modulates brainstem expiratory neuronal activity and delays the initiation of inspiration. In supine male subjects, we recorded ventilation, electrocardiogram, and blood pressure during 20-min epochs of baseline, slow-deep breathing, and recovery. In *in situ* rodent preparations, we recorded brainstem activity in response to pulses of perfusion pressure. We applied a well-established respiratory network model to interpret these data. In humans, the latency between HBs and onset of inspiration was consistent across different breathing patterns. In *in situ* preparations, a transient pressure pulse during expiration activated a subpopulation of expiratory neurons normally active during post-inspiration; thus, delaying the next inspiration. In the model, baroreceptor input to post-inspiratory neurons accounted for the effect. These studies are consistent with baroreflex activation modulating respiration through a pauci-synaptic circuit from baroreceptors to onset of inspiration.

## Introduction

Coupling of the respiratory and the cardiovascular systems is anatomical, physiological and reciprocal. Anatomically and physiologically, the respiratory and cardiac systems may interact for efficient gas exchange in that it decreases cardiac work (Ben-Tal, 2012; Ben-Tal *et al*., 2012). Although cardiorespiratory coupling (CRC) is reciprocal, the effect of respiration (*i*.*e*., the slower rhythm) on the cardiovascular system (*i*.*e*., the faster rhythm) is more apparent than the effect of the cardiovascular system on respiration. Indeed, the modulation of heart rate by respiration was one of the first physiologic system properties described (early 16^th^ century). Although referred as a respiratory sinus arrhythmia (RSA), it is not an arrhythmia but rather an increase in HR during inspiration followed by a decrease during expiration (Billman, 2011). Various mechanisms contribute to RSA. These include mechanical, as negative pleural pressure increases venous return, and neural, in that pre-ganglionic cardiac vagal neural activity is respiratory modulated with increased activity during expiration and thus, lowering HR.

The reciprocal manifestation of CRC, the cardiovascular system driving changes in the respiratory system, is cardio-ventilatory coupling (CVC). More specifically, CVC refers to the onset of inspiration occurring at a preferential latency following the last heartbeat (HB) in expiration (Tzeng *et al*., 2003; Friedman *et al*., 2012). The physiologic purpose of CVC was suggested to align breathing to the cardiac cycle and, thus, optimize RSA and make gas exchange more energy efficient (Galletly & Larsen, 1998). The “cardiac-trigger hypothesis” implicates baroreceptor input as a mechanism involved in the consistency of the latency observed between HB and the onset of inspiration. Basically, CVC depends on intact baroreceptors, so according to the cardiac-trigger hypothesis, the pulse pressure initiates an inspiration via baroreceptor activation (Bucher, 1963). However, the central neural substrate mediating this coupling remains undefined (Galletly & Larsen, 1997; Tzeng *et al*., 2007).

The literature supports the hypothesis that increases in blood pressure facilitate expiratory rather than inspiratory motor activity and preferentially modulate expiratory compared to inspiratory brainstem neural activity (Bishop, 1974; Grunstein *et al*., 1975; Lindsey *et al*., 1998). Respiratory rate decreased and the duration of expiration (TE) increased during these sustained increases in blood pressure (Bishop, 1974; Grunstein *et al*., 1975). Subsequent studies recorded brainstem neural activity during baroreceptor activation (Richter & Seller, 1975; Lindsey *et al*., 1998; Dick & Morris, 2004; Dick *et al*., 2005; Baekey *et al*., 2010). Richter and Seller (1975) recorded inspiratory (I) and expiratory (E) modulated activity intracellularly from the caudal medulla during pulsatile increases in arterial blood pressure. They reported that increases in carotid sinus pressure inhibited I-but failed to alter E-activity even though baroreceptor activation depolarized E-modulated neurons during periods of spontaneous hyperpolarization (Richter & Seller, 1975). Twenty-five years later Lindsey and colleagues returned to the question of how baroreceptor activation affecting respiratory-modulated neurons. They recorded neural activity from 221 respiratory modulated neurons in the ventral respiratory column (Lindsey *et al*., 1998) during gradually applied and sustained increases in blood pressure. Consistent with previous work (Richter & Seller, 1975), I-neurons largely decreased their activity during baroreceptor activation (aug-I neurons (n=61) 42.6% decreased vs. 8.2% increased; dec-I neurons (n=69) 17.4% decreased and 8.7% increased). Similar to I neurons, 22.9% of aug-E neurons (n=48) decreased rather than increased (14.6%) their activity during baroreceptor activation. In contrast, 32.6% of post-I neurons (n=43) increased and only 14% decreased their activity during the sustained baro-activation. Even though the majority of respiratory-modulated neurons did not change their firing frequency during the sustained baroactivation, increases in the post-I neural activity was associated with TE prolongation.

In a preliminary publication, we examined the effect of transient pressure pulses that inhibited sympathetic nerve activity and delayed the onset of the next inspiration on TE and on medullary neural activity *in situ* rodent preparations (Baekey *et al*., 2010). We reported an instance of two simultaneously recorded expiratory neurons, one with decrementing discharge pattern (post-I) and the other with augmenting activity (aug-E). When a short arterial pressure pulse was delivered during expiration, the post-I neuron increased and the aug-E neuron decreased their activities. These changes in activity were associated with a prolongation of TE. As the pulse subsided the post-I activity decreased and the aug-E neuron became reactivated. This was anecdotal evidence that the resetting of the respiratory rhythm was mediated primarily by the activation of the post-I activity.

The cardiac-trigger hypothesis implies that baroreceptor activation should initiate inspiration by activating pre-I activity. In contrast, published data indicates that baroreceptor activation affected E-modulated activity. Here, we expand our preliminary observations and test the hypothesis that CVC is mediated by the baroreceptors sensing the arterial pulse pressure and act by modulating post-I and expiratory neural activity (Baekey *et al*., 2008; Baekey *et al*., 2010). We explored this theoretical mechanism of CVC by using data from conscious humans, *in situ* rodent preparations, and mathematical modeling. We assessed the relationship between the HB and the onset of inspiration during normal and slow deep breathing in humans and during transient baroreceptor activation whilst recording brainstem respiratory neural activity in rodent *in situ* preparations. Then, from these rodent data, we developed a mathematical model of respiratory-baroreflex interaction and simulated human data to evaluate the possibility that the CVC may be due to the recruitment of expiratory neurons involved in determining the duration of expiration and the inspiratory onset.

## Results

### A hallmark of HB distribution is a preferred interval between the last HB during expiration and the onset of inspiration in human subjects (n=10, males)

CVC manifests itself as partial synchronization between HBs and respiratory oscillations and thus, is a property of CRC. However, due to substantial variability of respiratory phase durations respiratory phase-resetting due to this synchronization can be difficult to detect and characterize. Consistent with previously used approaches (Tzeng *et al*., 2003; Friedman *et al*., 2012), we found that the timing of the HB occurrence relative to the onset of the closest inspiratory period has the best defined structure compared to other metrics, e.g. HB phase within the respiratory cycle. Even though the latter metric appears similar, due to high variability of the respiratory cycle duration, the same time interval can result in drastically different phase difference.

In Figure 1, the raster plot has a pronounced structure indicating strong CVC for this individual. Each blue dot represents a HB; the *x*-coordinate is the time when the HB occurred, and the *y*-coordinate is the time interval between the HB and the onset of the closest inspiratory period. Negative times indicate that the HB occurred before the inspiratory onset, and positive times correspond to HBs that follow the closest inspiratory onset. Typical duration of the baseline respiratory cycle is about 5 seconds, so the *y* axis range covers approximately +/- half the cycle preceding and following each inspiratory onset. Specifically, the HBs immediately before and after E-to-I phase transition tend to occur at well-defined times (horizontal stripes of blue dots). The recording includes the three 20-min epochs (indicated by green to red vertical lines marking the beginning and end of each epoch, respectively): baseline (left), slow deep breathing (SDB, middle), and recovery (right). Interestingly, it is hard to see any difference in the structure of this plot between the three breathing epochs. One can notice, however, that the dots become less dense during SDB in the middle of the plot due to the smaller number of respiratory cycles.

**Figure 1.**
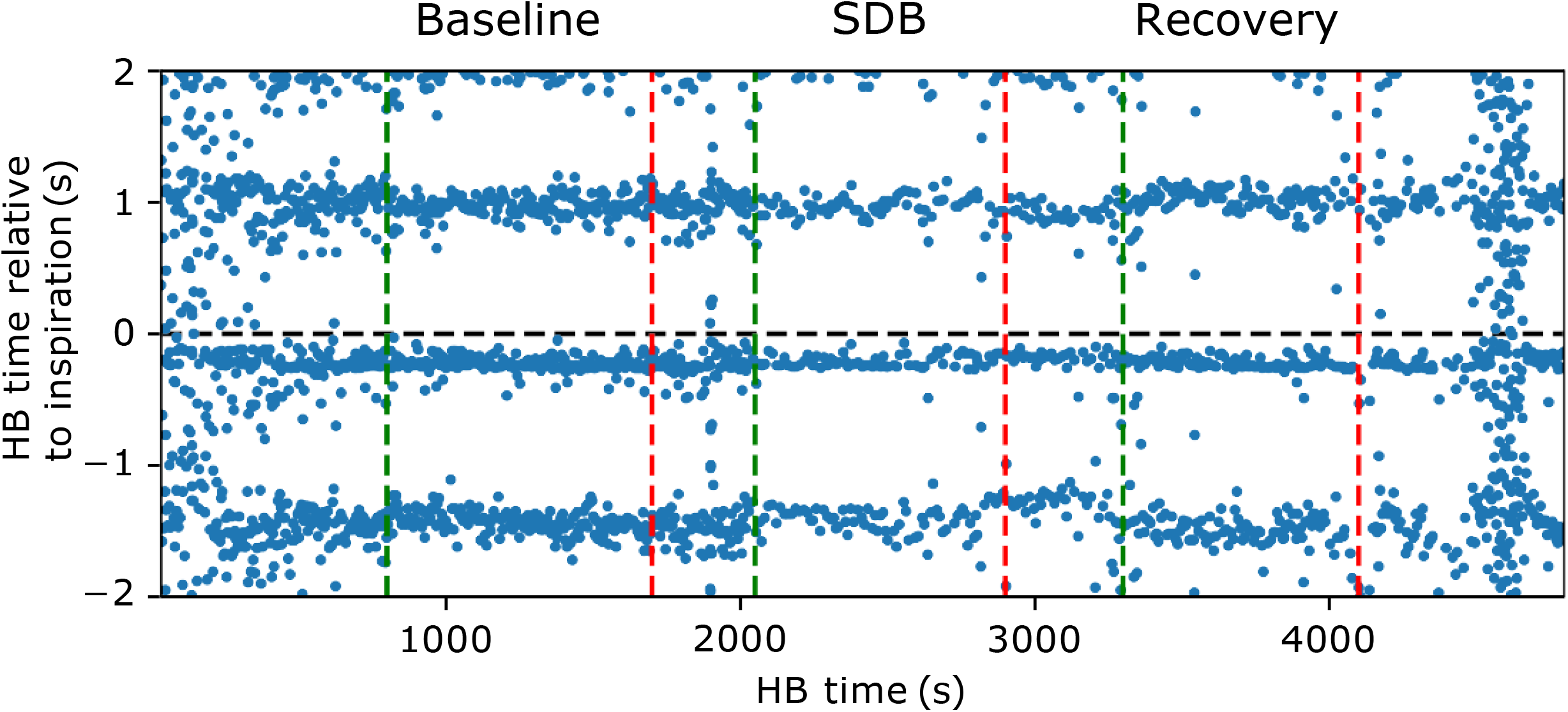
Temporal raster plot of the heartbeats relative to the inspiratory onset. Every blue dot represents a single heartbeat (representative data shown from one supine, male subject). The coordinates of the dot are the occurrence times in seconds of a heartbeat relative to the start of the recording (*x*-coordinate) and of the interval between the heartbeat time to the onset of inspiration of the closest breath (*y*-coordinate). Negative *y*-values correspond to heartbeats occurring at the end of expiration before the inspiratory onset, and positive *y*-values correspond to the heartbeats occurring after the inspiratory onset (shown by horizontal black dashed line). Vertical dashed lines show the beginning (green) and the end (red) of recording segments selected from the baseline, SDB and recovery parts of the experiment. We did not analyze the transition periods between baseline and SDB and between SDB and recovery.

We characterized this consistent CVC structure by estimating the HB probability distribution function as a normalized number of HBs occurring at a particular latency relative to the closest inspiratory onset for each experimental condition and each individual. This distribution is multimodal with each peak corresponding to a horizontal stripe of blue dots in Figure 1. Generally, the HB closest to 0 latency defined the narrowest horizontal stripe (this peak is shown in Fig. 2). The distance between peaks reflects the average RR interval of the corresponding individual. While the primary peak remains invariant, fluctuation in HR broadened secondary peaks (see Fig. 1). Based on these observations, we theorized that the CVC interaction is strongest between the HB closest to the inspiratory onset. In all individuals, the primary peak of the distribution of HR was within half a second before the inspiratory onset. Therefore, we focused on this window for Figure 2 and for further analysis.

**Figure 2.**
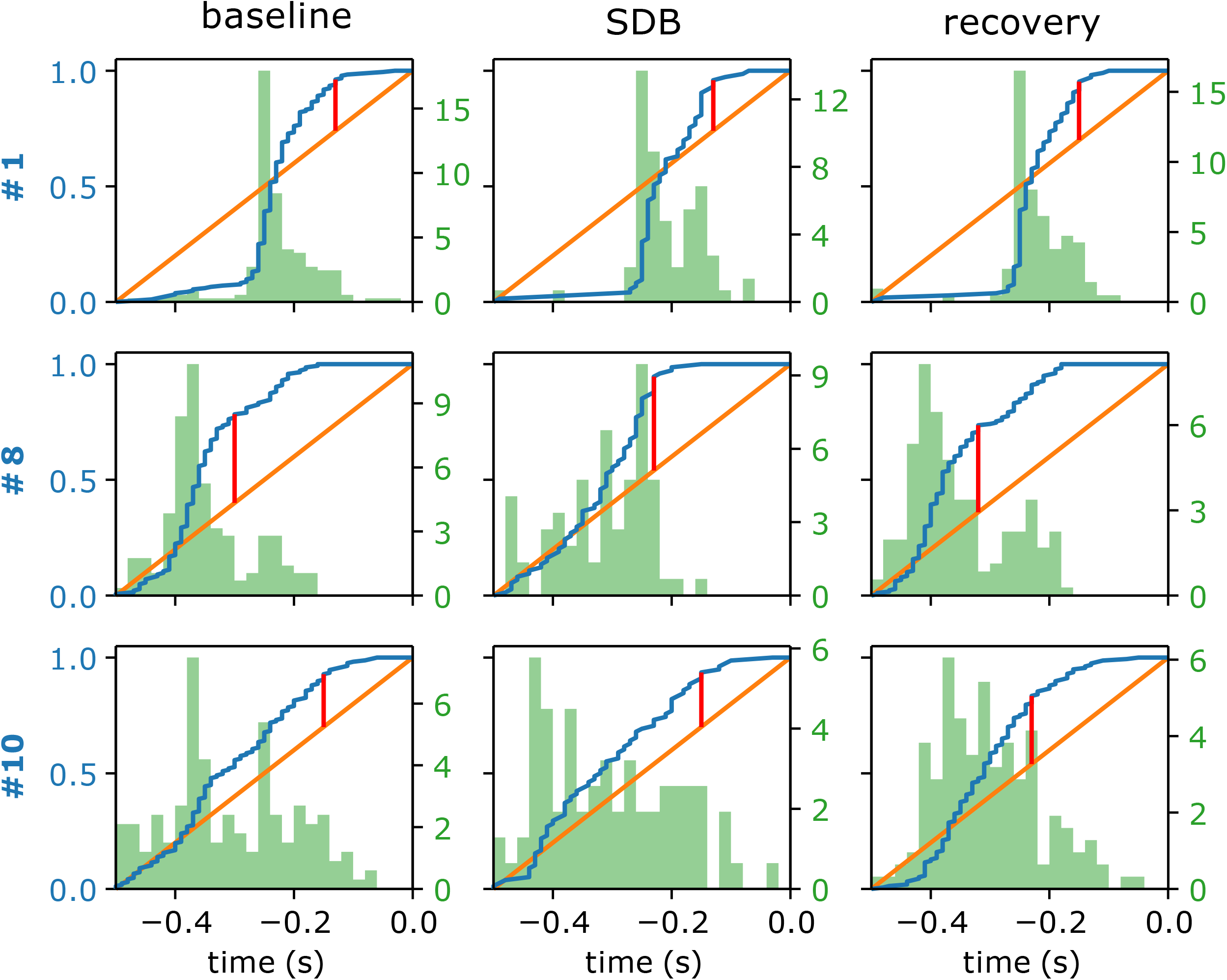
Distribution of heartbeats preceding the inspiratory onset. Each panel shows the cumulative distribution function (CDF, blue lines) as well as the histograms (probability density function, PDF, green bars) of the heartbeat occurrence times relative to the onset of the next inspiration (heartbeat latency) for the recording segments corresponding to baseline breathing, slow deep breathing (SDB) and recovery in supine male subjects. The three rows show data for three different individuals. Orange lines are the CDFs of the uniform probability distribution. Red bars indicate maximal distance between the actual CDF and the uniform CDF. The distributions for all 10 subjects were statistically significantly different from uniform distributions.

### Cardio-ventilatory coupling does not depend on the breathing pattern in human subjects

Figure 2 shows the estimated cumulative distribution functions (CDF, blue) as well as histograms (green) of HB latency before next inspiratory onset for three representative individuals during the three experimental conditions (all HBs for the ∼20 min periods of baseline, SDB, and recovery). If HBs and respiratory oscillations did not interact, then this latency would be distributed uniformly (Fig. 2, the orange line represents the uniform distribution). We quantified CVC as the maximal difference between the CDF and the uniform distribution (the red bars in Fig. 2). To evaluate statistical significance of these differences, we used the Kolmogorov-Smirnov statistical test which showed that 9 out of 10 individuals in our cohort had significant CVC (*i*.*e*., their latency distributions were significantly different from uniform at baseline). Furthermore, there was no significant difference between latency distributions obtained in different experimental conditions for a particular individual meaning that SDB in a relaxed state did not affect CVC.

Given that the difference between the actual and uniform latency distributions were significant, we used this difference as an index of coupling strength. Interestingly, this measure was consistent across the cohort (see Fig. 3A) as the standard deviation over the group is relatively small. Thus, consistent with our observation that the latency distribution does not depend on experimental condition for a particular individual, the group mean does not change significantly from baseline to SDB to recovery either (Fig. 3B).

**Figure 3.**
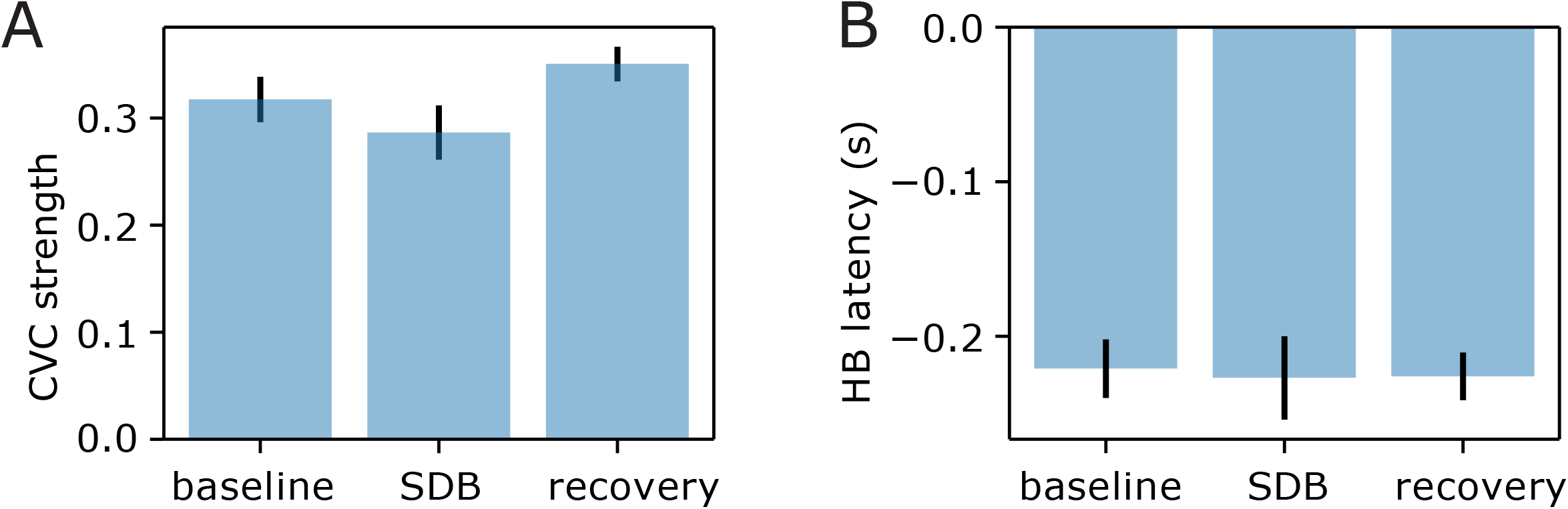
The measure of CVC strength and last heartbeat latency before inspiration. We used the maximal difference between the heartbeat latency cumulative distribution function (CDF) and the uniform distribution (see red bars in Fig. 2) as a measure of CVC strength in a particular individual. **A**. Group data for CVC strength which appeared to be consistent among individuals and did not vary significantly across the three experimental conditions. **B**. The characteristic heartbeat latency from inspiration (calculated as *x*-coordinates of the red bars in Fig. 2) also had similar values (approximately 200ms) across individuals and did not change significantly from baseline to SDB to recovery.

The structure of the latency distribution had a common characteristic feature; HBs were unlikely to occur during the short period of time (∼200 ms) before inspiratory onset (Fig. 2). Thus, the maximal positive difference between the CDF and the uniform distribution was observed right at the beginning of this period (see *x* coordinates of the red bars in Figure 2). We used the corresponding times to estimate the characteristic latency for each individual between the last HB and the onset of inspiration in each experimental condition (see group data in Figure 3B). We found that this latency was consistent across individuals and did not depend on experimental conditions.

### Pressure pulses delivered during expiration evoke a delay in the inspiratory onset in the rodent artificially perfused, brainstem preparation

As noted, the prevailing cardiac-trigger hypothesis is that arterial baroreceptors mediate the interaction between HBs and the respiratory rhythm generator. Baekey *et al*. (2010) correlated arterial pressure and respiratory activity in the artificially perfused brainstem preparation in rats (Fig. 4A). Specifically, isolated and solitary pressure pulses during the expiratory phase of the breathing cycle enhanced post-I activity, attenuated augmenting expiratory activity (aug-E) and delayed the onset of next inspiration (Fig. 4B).

**Figure 4.**
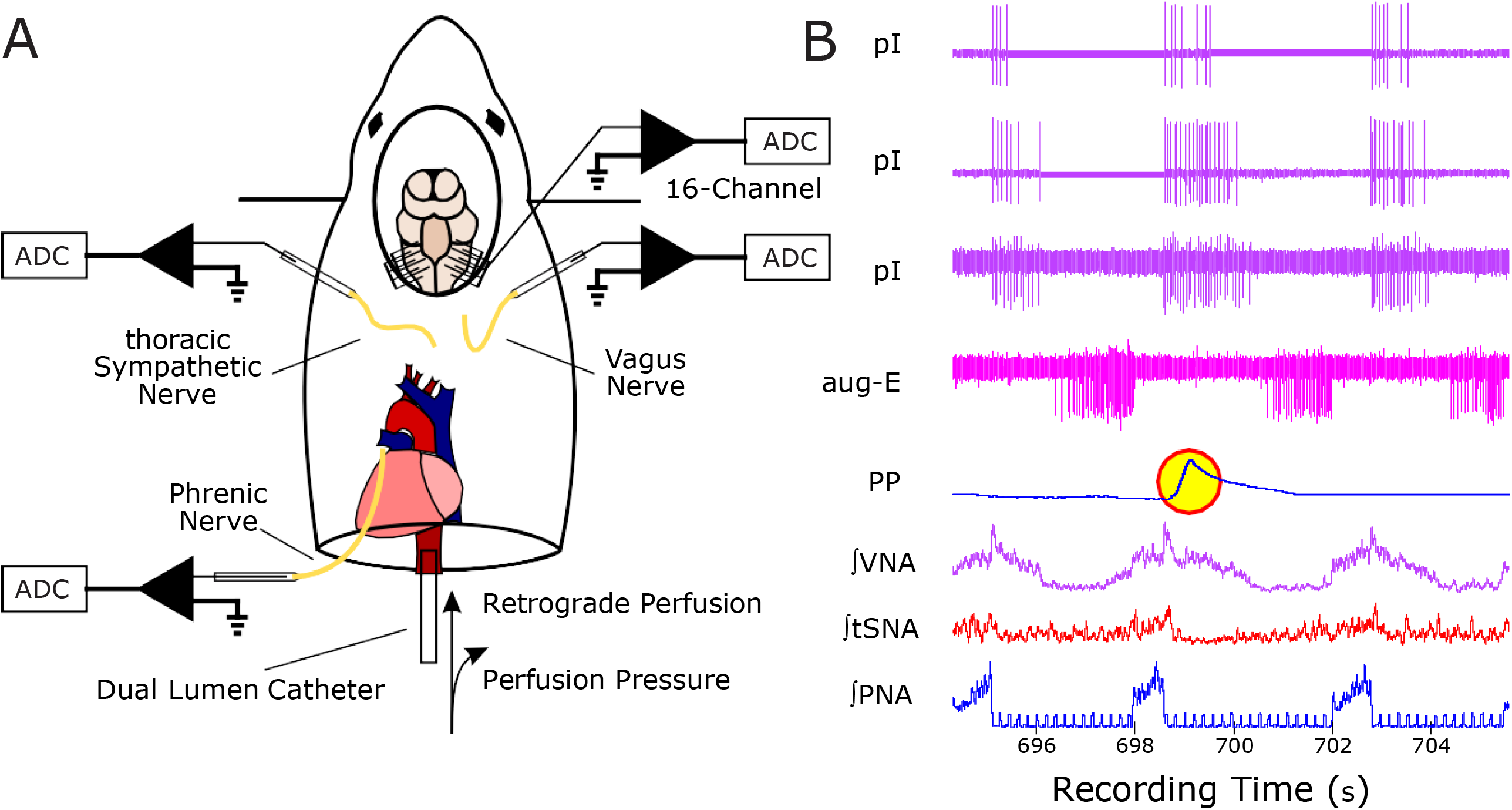
Experimental setup of artificially perfused brainstem-spinal cord preparation in a rodent. **A**. The preparation is referred to as *in situ* because the brainstem, spinal cord and connectivity to peripheral mechano-, baro- and chemosensory and to homeostatic motor fibers remain intact. Thus, reflex evoked responses can be recorded. **B**. Traces of the physiologic recordings. A pulse in the perfusion pressure (PP) can be delivered in different phases of the respiratory cycle defined by phrenic nerve activity (PNA, blue trace). Bursts in PNA correspond to the inspiratory phase and interburst intervals are expiratory phases. As shown in this example, when the pressure pulse occurs during expiration it noticeably delays the onset of the next inspiratory burst in PNA (i.e. prolongs expiration). It also causes a dip in thoracic sympathetic nerve activity (tSNA, red trace). Neural activity is recoded extracellularly by 16-channel multielectrode array. Examples of neuronal activity traces are shown in violet and pink. First three neurons exhibit post-inspiratory discharge pattern (pI) with stronger firing during the pressure pulse. In contrast, the fourth neuron (aug-E) that fires at the end of expiration, reduces its activity during perfusion pressure excursion.

We analyzed these data further (Fig. 5). The eupneic respiratory cycle consists of three phases: inspiration, post-inspiration and late expiration (E2), during each of these phases different populations of respiratory neurons are active. First, we confirmed that the effect of a single pressure pulse depended on the respiratory phase in which it was delivered. We found that pressure pulses during inspiration had no significant effect on inspiratory phase duration (−4.9 ± 1.7% (mean ± SE hereinafter), p = 0.055), while pulses during post-inspiration or E2 significantly prolonged expiratory duration: pulses delivered during post-inspiration increased time of expiration by 15.1 ± 2.4%, p < 0.001; pulses delivered during E2 phase increased time of expiration by 18.4 ± 3.6%, p = 0.008 (Fig. 5). Even though a significant difference was not apparent between pulses during post-I versus E2 phases, pulses during E2 phase tended to have a stronger effect on expiratory duration, which was consistent with Baekey *et al*. (2010) report.

**Figure 5.**
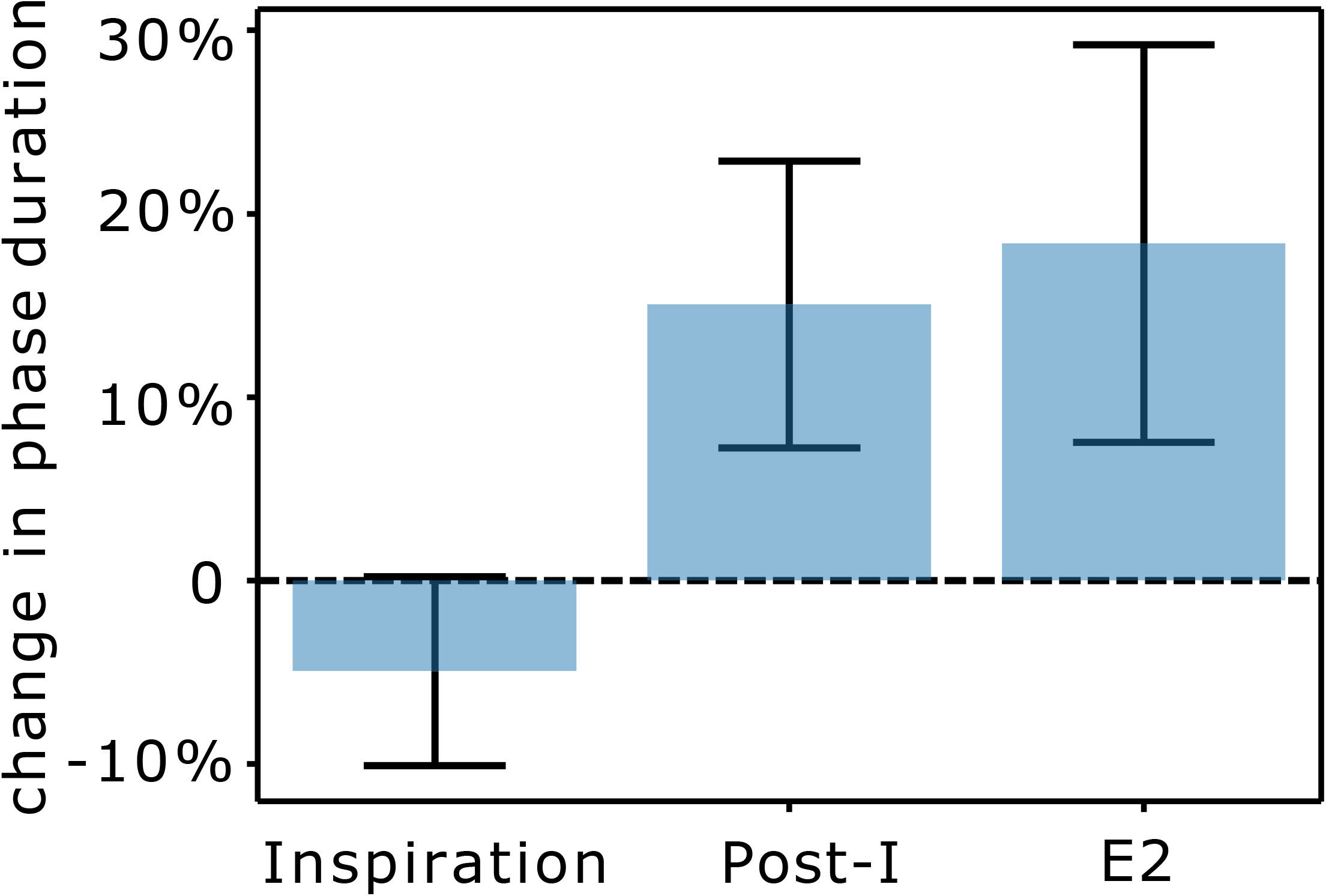
The effects of pressure pulses delivered in different phases of the respiratory cycle on the respiratory cycle duration. We determined the phase of pressure pulse from its peak. In the *in situ* preparation, if the pressure pulse occurred in inspiration (I, n=9), then it had no significant effect on cycle duration. But when delivered during first half (post-I, n=11) or the second half of expiration (E2, n=9), it prolonged the expiratory phase and thus increased cycle duration. Error bars represent the mean ± SD.

### During baroreceptor stimulation, post-inspiratory (post-I) neuron activity is increased, and augmenting expiratory (aug-E) neuron activity is decreased in the Bötzinger complex (BötC) of rodents

There are two major phenotypes of expiratory neurons. Post-I neurons, which are active in the first part of expiration have their maximal firing rate at the beginning of expiration and exhibit a decrementing firing pattern during expiration. Aug-E neurons fire after post-I neurons during expiration with an augmenting firing rate that terminate abruptly before the subsequent inspiration (see gray traces in Fig. 6A, B, upper panels). Baekey *et al*. (2010) reported a pair of simultaneously recorded post-I and aug-E during pressure pulses. The firing of the post-I neuron increased while the aug-E neuron decreased during the pressure pulse (see Figure 5 in (Baekey *et al*., 2010). To quantify this effect, we calculated the percent difference in average firing rate of post-I and aug-E neurons between unperturbed respiratory cycles and cycles with pressure pulses during expiration. We also calculated the same measure for the inspiratory neurons when the perturbation was delivered during inspiration (see group data in Fig. 6C). We found that inspiratory neurons did not change their firing rate significantly during pressure pulses coincident with inspiration (p = 0.95), which was consistent with no change in inspiratory duration when pulses arrived during inspiration. However, if the pulse occurs during expiration, post-I neurons increase their average firing rate by 27.8 ± 5.9% (p = 0.047) and aug-E neurons decrease their average firing rate by 39.2 ± 5.8% (p = 0.002). A closer look at their firing profiles showed that post-I neurons increase their firing right after the pressure pulse arrives (Fig. 6A) and continue firing until the pressure deviation ends, which coincides with the onset of the next inspiration. The effect on aug-E firing is largely opposite. These neurons reduce their firing right after the pressure pulse starts and then gradually come back as the pressure returns to its baseline (Fig. 6B).

**Figure 6.**
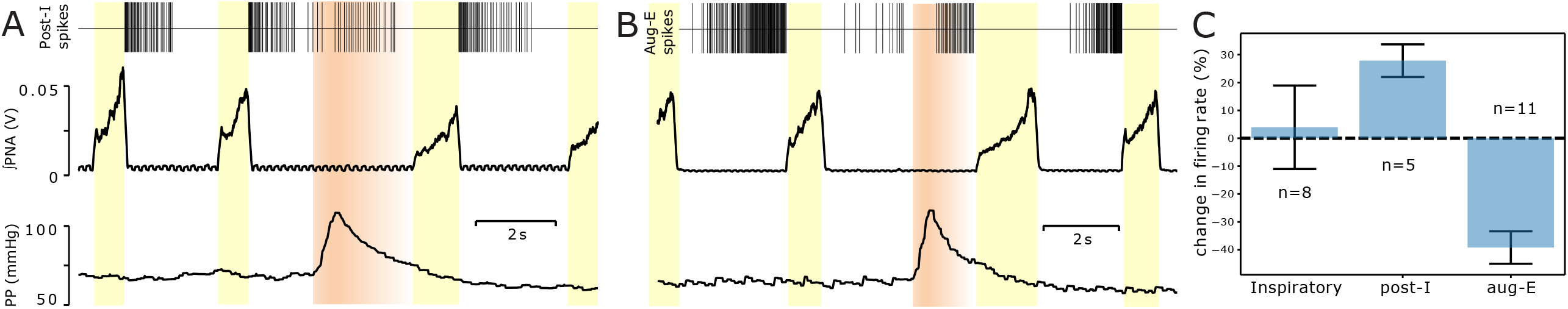
Effects of pressure pulses on firing of expiratory-modulated brainstem neurons. **A**&**B**. Tracings from top: representative post-I neuron (**A**) and aug-E neuron (**B**), perfusion pressure (red) and integrated PNA (black). Gray thick curve in the top panel represents the cycle-triggered average of the firing rate of these neurons in unperturbed cycles. The pressure pulse was delivered at the time when the post-I neuron (**A**) would cease firing and when the aug-E neuron (**B**) would be augmenting. During baroreceptor stimulation induced by the transient pulse pressure, the firing rate of the post-I increased (**A)** whereas the aug-E neuron decreased then recovered the perfusion pressure decreased (**B**). **C**. Group data summarizing the effect of pressure pulses on the activity of neurons of different firing phenotypes (I (n=8), post-I (n=5), and aug-E (n=14)). When the pulse was delivered during inspiration, it had no significant effect on the average firing rate of the recorded inspiratory neurons. When the pulse was delivered during expiration, we registered significant increases in post-I neurons activity and decreases in aug-E activity.

### Mathematical modeling of baroreceptor input to post-I neurons of the respiratory central pattern generator (CPG) explains the effects of baro-stimulation in rodents

Baekey *et al*. (2010) hypothesized that the delay in inspiratory onset induced by perfusion pressure pulses was mediated by arterial baroreceptor input to the brainstem expiratory neurons. They demonstrated the plausibility of this hypothesis by computational modeling, that second-order baroreceptor neurons in the *nucleus tractus solitarii* (nTS) send excitatory projections to the post-I neurons in the BötC. We implemented the same idea in a simpler and more mathematically tractable rate-based model of the respiratory CPG first published by Rubin *et al*. (2009) and then tested by other researchers against a large number of experimental findings, e.g. (Rubin *et al*., 2011; Molkov *et al*., 2014; Ausborn *et al*., 2018). Populations of neurons of the same phenotype in this model are described in terms of population firing rate, which is a predefined monotonous function of average membrane potential over the population (see Methods). Membrane potential dynamics follows conductance-based description similar to typically used in Hodgkin-Huxley-like formalism but with spiking currents excluded. We assumed that the conductances of excitatory and inhibitory synaptic channels depended linearly on the firing rates of projecting presynaptic populations with coefficients representing synaptic weights of the corresponding connections.

According to Smith *et al*. (2007) whose work was a basis for the model by Rubin *et al*. (2009), the apneic respiratory pattern is a result of intrinsic dynamical properties of, and synaptic interactions between, neurons in preBötC and BötC of medulla oblongata (Fig. 7). PreBötC mostly contains inspiratory neurons (*i*.*e*. those that fire during inspiration) and BötC mostly contains expiratory neurons. There is a large population of excitatory recurrently connected inspiratory neurons in preBötC which is capable of endogenous rhythm generation in isolation (Smith *et al*., 1991). Endogenous bursting depends on the expression of persistent sodium current in a subpopulation of these neurons (Koizumi & Smith, 2008). The expiratory neurons are represented, as noted, by two mutually inhibiting populations with post-I and aug-E discharge patterns. They also reciprocally interact with an inhibitory population of preBötC inspiratory neurons labelled early-I in Fig. 7.

**Figure 7.**
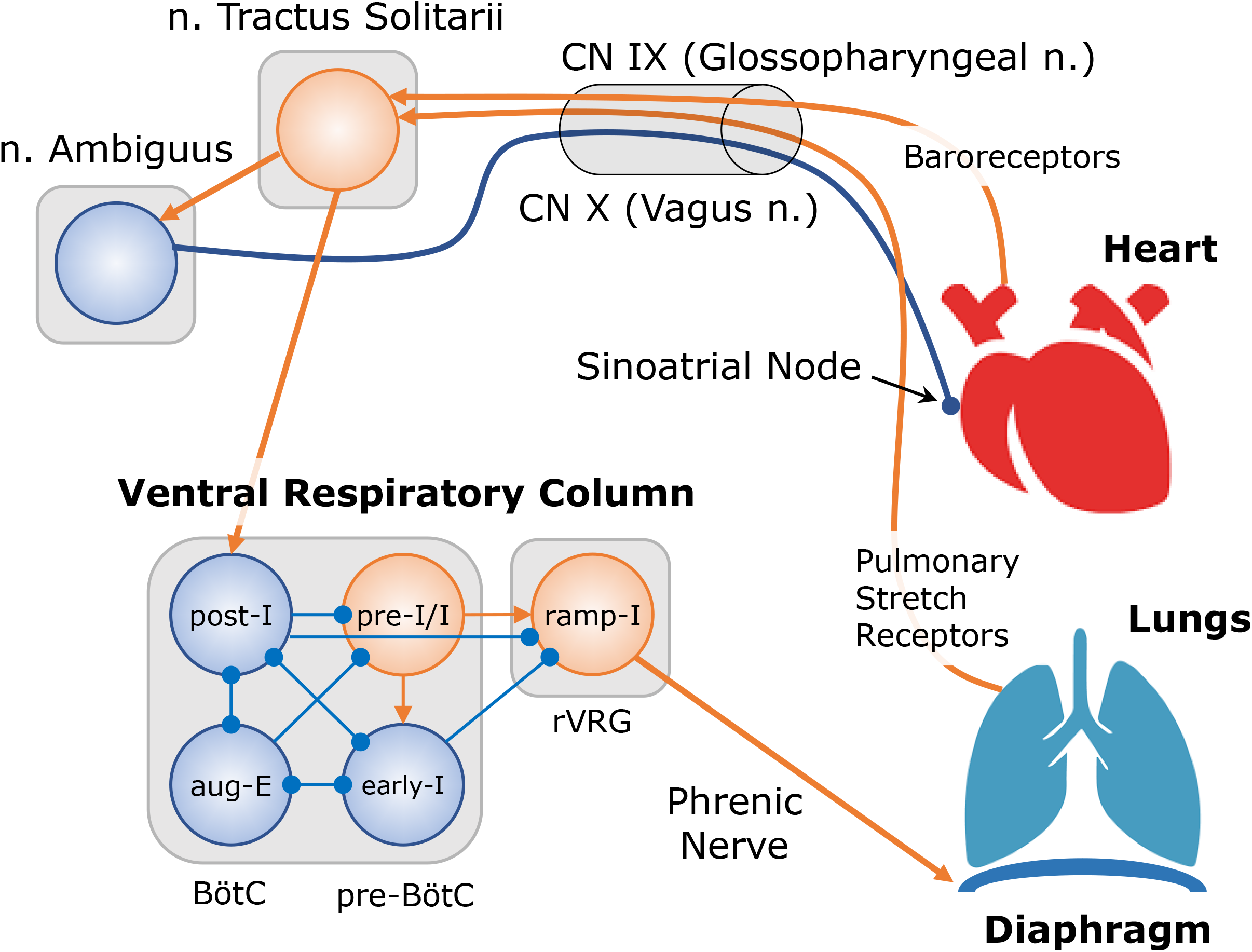
Model schematic for cardio-respiratory interactions. The respiratory central pattern generator (CPG) is represented by interconnected populations of neurons in Bötzinger (BötC) and pre-Bötzinger (pre-BotC) complexes that contribute to the activity of the phrenic premotor population (ramp-I) in the rostral ventral respiratory group (rVRG). These neurons define the activity of the diaphragm and lung inflation. In the absence of ramp-I activity, the lungs passively deflate. The lung volume is decoded by pulmonary stretch receptors that send synaptic inputs to the pump cells located within the nucleus tractus solitarius (nTS) through the vagus nerve. Pump cells excite BötC post-I neurons which creates a negative feedback loop for off-switching inspiratory activity (Hering-Breuer reflex). Nucleus Ambiguus contains a population of cardiac neurons that modulate heart rate by inhibitory inputs to the sinoatrial node. The cardiac neurons receive inputs from respiratory populations and/or pump cells so that their output becomes respiratory modulated and thus serves as a mechanism for respiratory sinus arrhythmia and blood pressure oscillations (Traube-Hering waves) in the model. Arterial baroreceptors encode the blood pressure value and send this information to the nTS second-order neurons in the baroreflex arc. The latter project to the post-I neurons in the BötC thus creating a beat-by-beat arterial pressure input to the respiratory CPG underlying cardio-ventilatory coupling. Through these mechanisms the heartbeat can affect the timing of the next breath.

The succession of respiratory phases in the model occurs as follows (see (Smith *et al*., 2007) and (Rubin *et al*., 2009) for mechanistic and mathematical details, respectively). Post-I and early-I neuronal populations form a half-center oscillator based on their mutual inhibition and spike-frequency adaptation properties. Due to the latter, both have decrementing activity patterns (Fig. 8). Adaptation of post-I firing disinhibits aug-E neuron activity, which emerges at some point of expiratory phase and then gradually increases towards the end of expiration. In our extension of the model, we assumed that the pressure pulse induces a baroreceptor activity profile as shown by the bottom trace in Fig. 8. This profile was used as a direct excitatory synaptic input to the post-I population (red arrow from nTS to post-I in Fig. 7). So, for the duration of this input the firing rate of the post-I population was increased which led to a delayed transition from expiration to inspiration in that respiratory cycle (see dashed traces in Fig. 8). Besides, due to increased post-I activity during baroreceptor activity pulse, aug-E activity was depressed.

**Figure 8.**
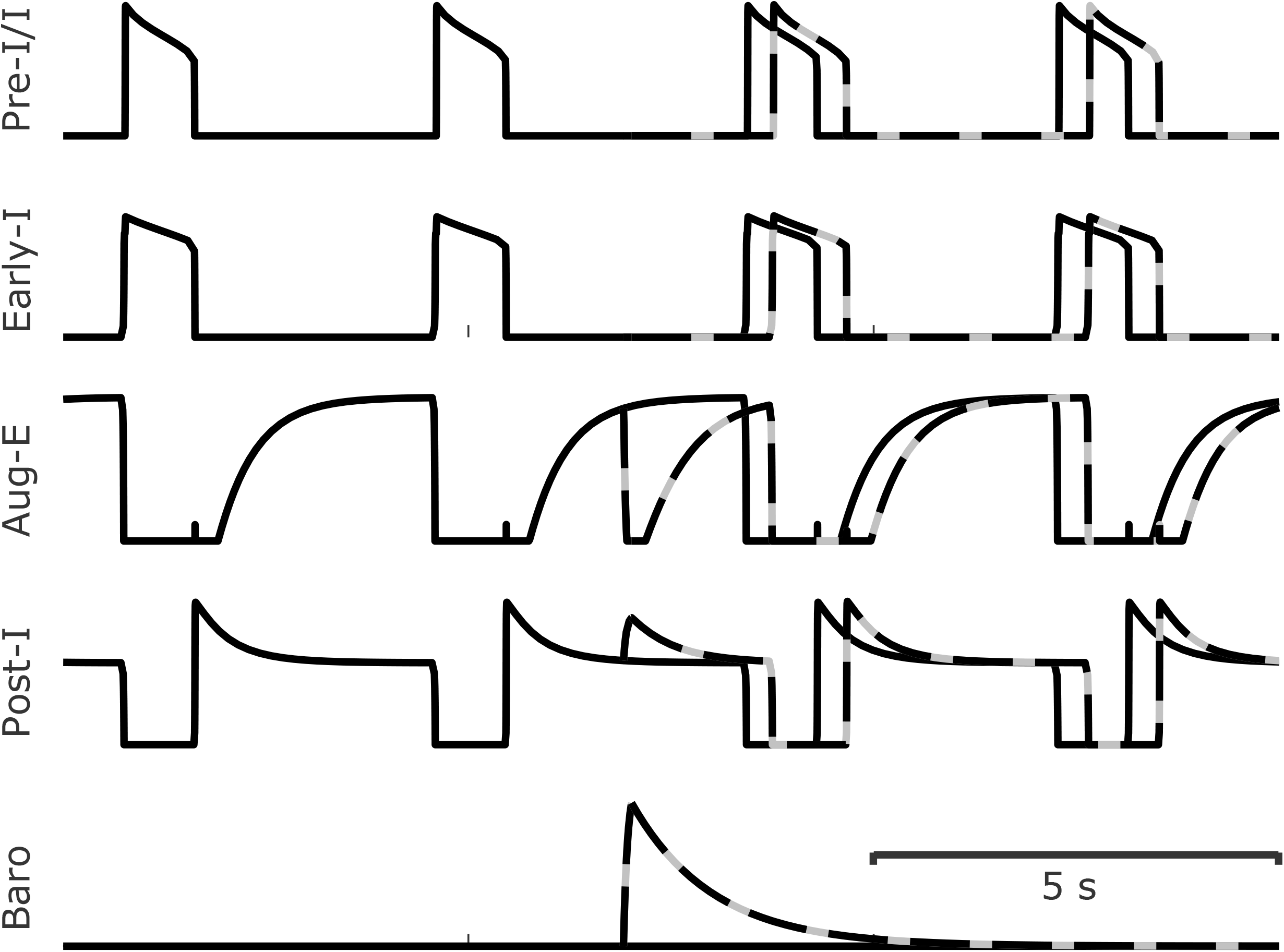
Modeling the effect of transient baroreceptor stimulation. Traces from the top: activity of four neuronal populations representing the respiratory CPG: pre-I/I, early-I, aug-E and post-I (see Fig. 7). Black traces represent unperturbed activity. After inspiration ends, the post-I population activates strongly and then adapts, gradually releasing the aug-E population from inhibition. This post-I adaptation eventually allows the inspiratory populations (early-I and pre-I/I) to activate. Dashed traces show the CPG activity in the presence of a baroreceptor stimulus (bottom trace). When it arrives centrally, the post-I population reactivates again, inhibits the aug-E population and prevents inspiration from starting until the baroreceptor activity wanes.

In summary, the model explains the delay in the inspiratory onset after baro-stimulation as well as the changes in neuronal discharge patterns by excitatory synaptic inputs from nTS baroreceptor neurons to the post-I population of the respiratory CPG. Importantly, as long as the pulse arrives late enough in expiration, the duration of the baroreceptor activity pulse defines the duration of the delay.

### Mathematical modeling of repetitive baroreceptor input to post-I neurons from HB-produced pressure pulses creates a HB distribution similar to human data

We implemented a cardio-respiratory mathematical model that includes several interaction mechanisms between the two systems. The model is based on our previous work (Molkov *et al*., 2013; Molkov *et al*., 2014) where we incorporated feedback (also known as Hering-Breuer reflex) from pulmonary stretch receptors to the central respiratory neural circuits. We included a simple model of the heart to generate HB times as a Poisson process with the rate modulated by the respiratory activity. We engineered this modulation to produce CRC/RSA consistent with the one we published based on the analysis of the same experimental group (Barnett *et al*., 2020). We borrowed the model of arterial pressure dynamics from the same publication, which shows pulsatile dynamics of the pressure with slow respiratory modulation (Traube-Hering waves). Finally, we included a simple model of baroreceptor activity as a signal proportional to the excess of arterial pressure over a certain threshold and used this signal as an excitatory input to the post-I neurons of the respiratory CPG. We used the synaptic weight of this input as an independent parameter that defines the CVC gain. Figure 9 shows the dynamics of the main physiological outputs simulated by the model in comparison with their experimental counterparts for both baseline and SDB conditions.

**Figure 9.**
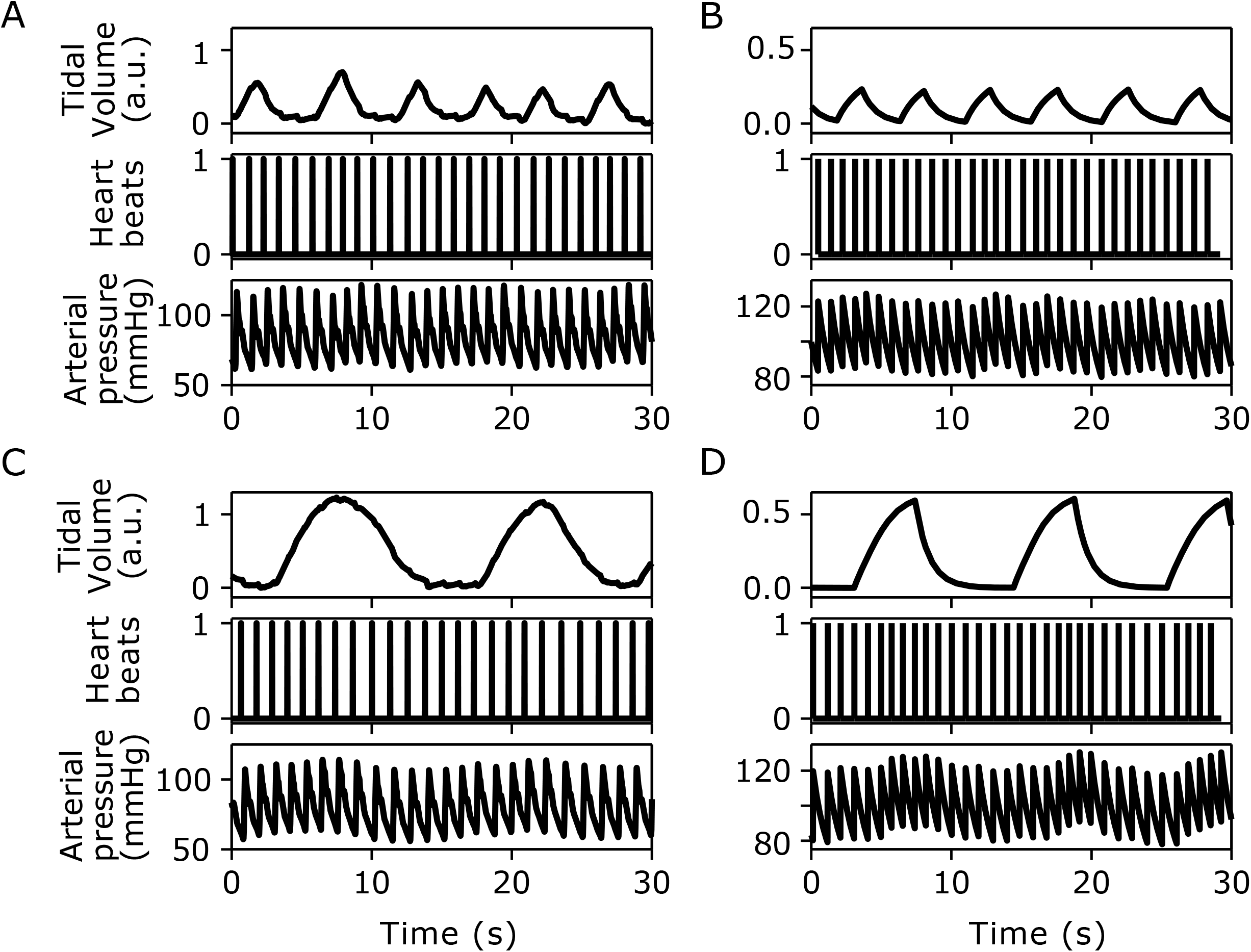
Representative data from a subject (A&C) compared to model simulations (B&D). **A&C**. Traces (30 sec) of tidal volume, time stamp for ECG R-peaks and brachial intra-arterial pressure during baseline (A) and slow deep breathing (SDB, C). **B&D**. Dynamics of the corresponding variables in the model mimicking baseline (B) and SDB (D) conditions (see text). We tuned the model in such a way that respiratory phase durations, HR, systolic and diastolic pressures as well as variabilities of all these metrics in both baseline and SDB conditions matched our experimental group data we previously published (Barnett *et al*., 2020); see Fig. 9. We varied the CVC gain in the model to determine the range in which the model demonstrated heartbeat distributions similar to the experimentally observed ones (Fig. 2).

In Fig. 10A, we present simulation results for the HB distribution in the same format as the human data in Figure 2. These simulations had the same durations as experimental recordings. For all three conditions (baseline, SDB and recovery) HB distributions exhibit a range of latencies immediately preceding the onset of inspiration (0) where HBs were unlikely to occur. The characteristic latency between the last HB and the onset of inspiration (calculated in the same way as we did for data; see above) was about 200 ms for simulations of baseline conditions. Interestingly, the characteristic latency during simulated SDB was slightly longer (∼300ms).

**Figure 10.**
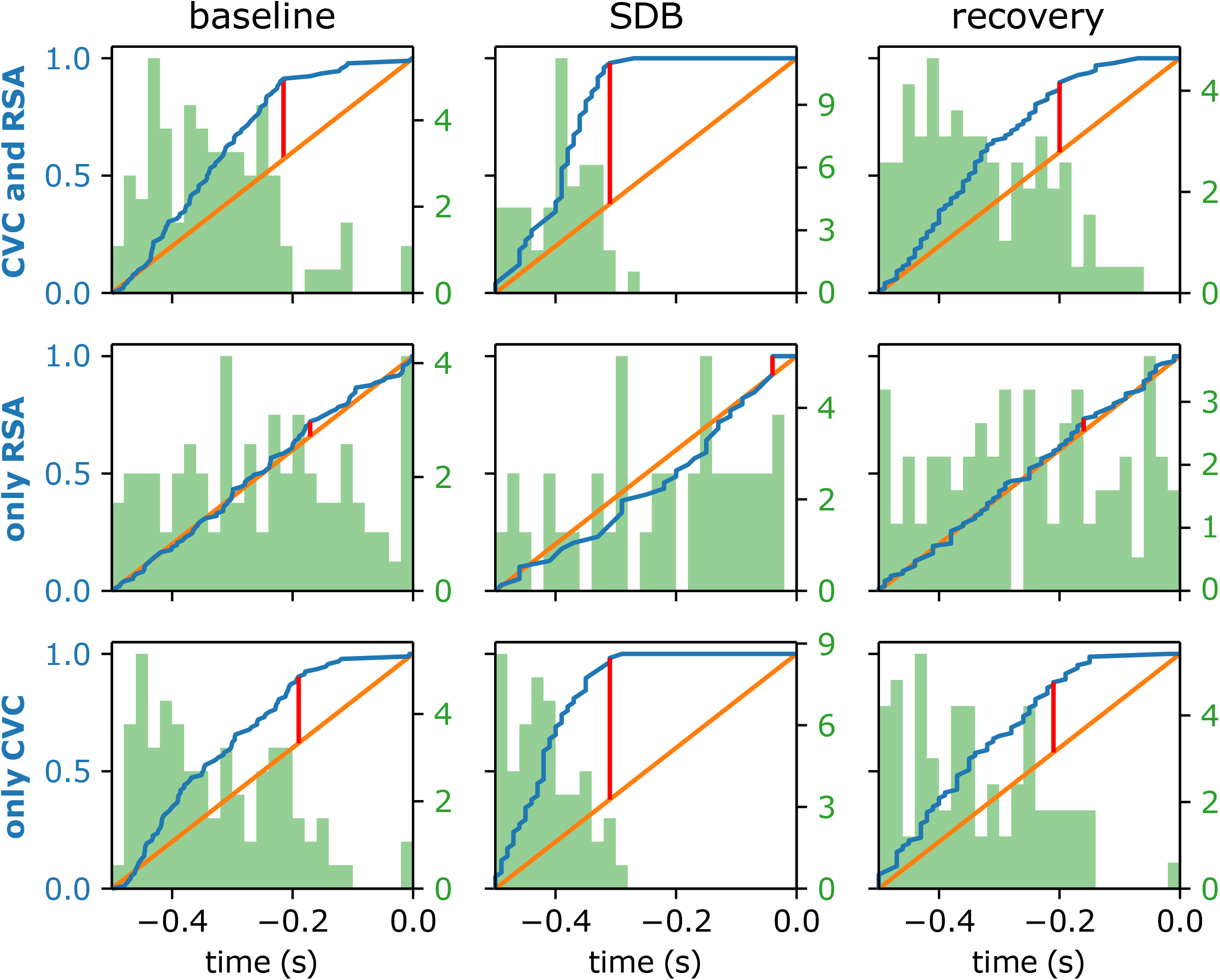
The model qualitatively reproduces the heartbeat latency distributions as long as baroreceptor-to-respiratory network pathway is functional. Each panel shows the cumulative distribution function (CDF) of heartbeat latency before inspiration (blue lines) and corresponding probability density function (PDF) histograms (green bars) in the 0.5s time interval preceding the inspiratory onset. Orange lines depict CDFs of uniform distributions. Red bars indicate maximal differences between the heartbeat latency CDFs from the model and uniform distributions. For the first row of the plots, the model included interactions between respiratory and cardiac systems in both directions, i.e. RSA and CVC. Both conditions, baseline and SDB in human subjects, feature a 200-300ms gap in heartbeat latency distributions. In the second row, we disrupted the CVC by setting the NTS-to-post-I synaptic weight to 0 (see Fig. 7), which made the heartbeat latency distribution statistically indistinguishable from the uniform distribution (orange lines). The third row was constructed by removing the respiratory modulation of NA cardiac neurons (underlying RSA, see Fig. 7) from the model while retaining CVC, which did not have a significant effect on the distributions (compare with the first and the third rows).

Our model had two coupling mechanisms between respiratory and heart rhythms, we evaluated their individual contributions to the CVC phenomenon in the model. First, we removed the baroreceptor input to the post-I neurons of the respiratory CPG and repeated the same simulations. We found that without this input the distribution of HBs becomes statistically indistinguishable from the uniform distribution (Fig. 10B) meaning that there is no CVC. In contrast, CVC remained after removing respiratory modulation of the heart rate (RSA) so RSA did not affect the distribution of HBs relative to inspiratory onset. (Fig. 10C).

## Discussion

The coupling of the respiratory and the cardiovascular systems is observed in a number of physiological scenarios. One key manifestation of such coupling is CVC in which a HB is more likely (or unlikely) to occur during certain phases of the respiratory cycle. The present study examines a neural mechanism by which the cardiovascular system can affect the respiratory pattern. Using a combination of human data, *in situ* animal data and mathematical modeling, we test the hypothesis that systolic peak pressure activates arterial baroreceptors and initiates CVC. This aspect of CVC is consistent with the cardiac-trigger hypothesis. However, when triggered, the baroreceptor afferent input through a pauci-synaptic neural pathway to the respiratory CPG delays the onset of the inspiration by activating the expiratory neurons. Thus, the underlying neural mechanism of CVC differs from that alluded to by the cardiac-trigger hypothesis in that it delays rather than initiates a phrenic burst.

In supine resting humans, SDB strengthens the magnitude of RSA and Traube-Hering waves (Dick *et al*., 2014b; Barnett *et al*., 2020), but CVC remains robust and unaffected. Thus, CVC is determined by an independent mechanism from that of RSA and TH waves and distinct from that of respiratory modulation of autonomic activity. There are several key experimental observations supporting the concept that baroreceptor responsiveness to blood pressure fluctuations mediate CVC.

First, transient increases in blood or perfusion pressure *in vivo* and *in situ* evoke an expiratory facilitatory reflex with the recruitment of expiratory motor activity and an increase in the duration of expiration (Bishop, 1974; Baekey *et al*., 2008; Baekey *et al*., 2010). The evoked cardio-sympatho-respiratory response depends on the respiratory phase (Baekey *et al*., 2008). Thoracic sympathetic activity and HR decreased whether the pulse was delivered in inspiration, post-inspiration or E2 phases. The magnitude of the autonomic decreases was respiratory phase dependent with the least effect occurring during inspiration, and the greatest in post-inspiration. While duration of expiration was not prolonged when the pulse was delivered in inspiration, it was prolonged following pressure pulses delivered in the post-I and E2 phases. The evoked response also depended on the magnitude of the pressure pulse. A pressure pulse as weak as 18 mmHg (or ∼25% above the mean perfusion pressure) evoked sympatho-respiratory response, in which sympathetic activity was inhibited and the TE was prolonged but the decrease in HR was minimal (Baekey *et al*., 2008).

Second, arterial pressure pulses resulting from the HB do modulate respiratory neuronal activity. Our previous studies in cats indicate that pulse pressure modulated expiratory activity recorded from isolated single brainstem neurons (Dick & Morris, 2004; Dick *et al*., 2005). Thus, the same neuron can express respiratory and pulse modulated activity. Once we realized this, it became an issue of identifying if this was simply a cardio-sympathetic control neuron expressing respiratory modulation or a respiratory neuron being baro-modulated. To resolve this, we characterized the axonal projection of the recorded neurons and identified recurrent laryngeal motoneurons and excitatory premotor inputs to the recurrent laryngeal motoneuronal pool (Dick *et al*., 2005). In these subgroups, we found that expiratory activity was preferentially affected by baroreceptor inputs and that activity could be facilitated or inhibited following a HB.

Third, breathing pattern variability depends on baroreceptor input. We (Dick *et al*., 2014a) and others (Galletly & Larsen, 1999) have noted that baroreceptor input increases ventilatory variability. In our study in anesthetized rats, variability of the respiratory frequency differed depending on whether the rodents had been conditioned to either chronic intermittent or sustained hypoxia (Dick *et al*., 2014a). After chronic intermittent hypoxic conditioning, CRC was strong and had minimal respiratory frequency variability, whereas after chronic sustained hypoxic conditioning CRC was weak and respiratory frequency variability greater. Surprisingly, this high respiratory frequency variability depended on the aortic depressor and carotid sinus nerves being intact (Dick *et al*., 2014a). In anesthetized humans, four types of coupling patterns occurred. Variability in respiratory frequency was lowest when the HB had a consistent number of beats, generally 3 or 4 beats, per breath. In contrast, coupling patterns in which the number of beats per breath varied resulted in a variable respiratory frequency. The respiratory cycle duration would transition or oscillate and maintain an integer relationship for HBs per breath (Galletly & Larsen, 1999).

### Limitations and future directions

Tonic arterial baroreflex afferent activity is modified acutely throughout the cardiac cycle and HB to HB. The arterial baroreceptors are most active during the rising phase of arterial pressure with each HB and their activity is dynamic in relation to blood pressure. The change in arterial pressure is a key determinant of tonic activity in the baroreceptor neurons. The rate of pressure change, the duration of the pulse, prolonged changes in pressure, and baroreceptor adaptation are all related to changing central baroreflex afferent input. Therefore, it is essential that we consider that a longer pressure pulse (such as delivered in the *in situ* experiments in this paper) or an overall shift in mean blood pressure during exercise or hypertension in humans may indeed reveal different results than the transient baro-activation over the course of the systolic phase of a single cardiac cycle. This is a recognized limitation of the study and supports additional studies to explore differences, if any, resulting from the use of the baroreflex activation pattern with the width of a single cardiac cycle versus a longer period of activation. For example, the use of lower body positive pressure is a longer baroreflex challenge than is brief neck pressure. Therefore, interpretation of those studies as it pertains to the relationship between baroreflex and ventilatory control should consider the nature of the afferent input under different pressure profiles. Also studies in humans that involve a sustained increase in arterial pressure or baroreflex gain resetting–such as intense exercise or studies in hypertensive patients–would be useful to better characterize CVC.

A phenomenon that inspired this study was CVC, manifested by a well-defined latency between the last HB during expiration and the inspiratory onset (Fig. 1). This latency did not depend on experimental conditions, *i*.*e*. normal vs. slow deep breathing, although the participants were relaxed and calm in both conditions (Fig. 3). In our simulations we observed that the latency increased as we reduced the frequency in the model to mimic SDB. We simulated SDB by reducing excitatory drive to key respiratory populations thus decreasing their excitability. We theorize that the stereotypic latency between the last HB and inspiratory phase is caused by the transient baroreceptor input to expiratory neurons and that the profile of this input is dictated by the arterial pressure, which gradually relaxes between HBs. The increased latency for the expiratory-to-inspiratory transition occurs in the model due to reduced excitability of simulated neuronal populations while the strength of the external baroreceptor input remained the same. Therefore, the expiratory-to-inspiratory transition required the simulated blood pressure to fall to lower levels during SDB compared to normal breathing. This, however, implies a specific control mechanism that participants employ to implement slow deep breathing pattern.

Experimentally, the latency between the last HB and inspiration was invariant during normal and SDB suggesting that our mathematical implementation of breathing control had flaws. In this study, we focused on a coupling mechanism between cardiovascular and respiratory systems and used a popular model for respiratory rhythm generation. It would be interesting in the future, however, to explore whether other respiratory control mechanisms are compatible with the latency invariance above.

### Relevance

The direct coupling of inspiratory onset control to the cardiovascular system has important functional consequences. Inspiration facilitates the “respiratory pump” and can maintain stroke volume during hypovolemia (Skytioti *et al*., 2018). Convertino (2019) summarized the importance of the changes in intrathoracic pressure during inspiration to facilitate the respiratory pump in a range of hypovolemic conditions including hemorrhage and orthostatic hypotension. As such, rapid communication between the arterial baroreceptors and inspiratory control would be advantageous. Indeed, using several approaches, it has been reported that there is a relationship between blood pressure and ventilation with low pressure associated with high ventilation and vice versa. The later observation is aligned with our conclusion that baroreflex activation with high pressure delays inspiratory onset.

Lower body negative pressure, neck suction, and brief infusions of vasoactive drugs acutely alter baroreflex activity through transient changes in blood pressure. These techniques offer insight into the reciprocal interactions between arterial baroreflex and ventilatory control in humans. During lower body negative pressure, which unloads the carotid baroreceptors as blood volume is shifted to the lower limbs, ventilation and the respiratory pump are greatly increased (Koehle *et al*., 2010). Lower body positive pressure, which activates the baroreceptors as central blood volume and stroke volume increase, does not result in changes in ventilation. Our studies support that baroreceptor stimulation delays inspiratory onset; however, in the longer-term steady state increase in pressure generated by lower body positive pressure ventilation rate is unchanged. The duration of the baroreflex triggering and a baroreflex resetting to prolonged activation should be considered.

Further, pharmacological interventions aimed at changes in blood pressure alter ventilatory rate with increased blood pressure decreasing ventilation and vice versa (Stewart *et al*., 2011). There is a striking stimulation of ventilation, in particular of tidal volume, during rapid pharmacological infusion of vasoactive drugs (Oxford Maneuver) in human subjects. This so-called “ventilatory baroreflex” is not related to chemoreflex and the mechanisms are still under investigation (Stewart *et al*., 2011).

While generally considered to be baroreflex mediated, the above interventions may also have interaction with the chemoreflex. For example, hypoxic ventilatory response mediated by the peripheral chemoreceptors is increased during lower body negative pressure (Koehle *et al*., 2010). During severe hemorrhage combined with hypoxia there is activation of the peripheral chemoreceptors in the carotid body but this does not occur with low blood volume alone (Kumar & Prabhakar, 2012). That being said, the hyperventilation triggered in low volume states is associated with hypocapnia which is not a stimulus to the central chemoreceptors. Low volume alone did not result in activation of the peripheral chemoreceptors unless combined with low oxygen.

### Conclusion

Using a combination of animal data, human data and mathematical modeling, we explored the underlying mechanisms of CVC. We hypothesized that the HB derived pressure pulses entrained the respiratory pattern via baroreceptor mediated modulation of the initiation of inspiration. As each HB triggers blood pressure pulses and baroreceptor activation, a neural pathway that inhibits an inspiration is activated thus affecting the timing of the inspiration onset. If correct, it would be likely that the latency between a HB directly preceding inspiration and the inspiratory onset would depend on the duration of baroreceptor activity pulse and the transmission time to the ventral respiratory column. This hypothesis was further tested by using an SDB protocol in human subjects to probe if breathing can alter the linkage of HB to inspiration, and using an *in situ* brainstem-heart rodent model preparation in which pressure pulses can be introduced during different phases of respiration to test how the timing of the next inspiration is affected when the timing of baroreflex input to the brainstem is changed. We conclude that baroreflex activation modulates inspiration timing through a pauci-synaptic circuit from the baroreceptors to the ventral medullary respiratory column. Specifically, a transient pressure pulse during expiration increased post-I neuronal activity, decreased aug-E activity transiently, and delayed the next inspiration. The model supported the notion that baroreceptor input to post-I neurons accounted for CVC.

In summary, key findings of this study are: 1) In the human subjects, there was a stereotypic latency (∼200 ms) from the last HB during expiration to the onset of inspiration in both involuntary and voluntary breathing. The latency was unaltered during SDB; 2) In the rodent preparation, triggering of baroreflex input via an experimental pressure pulse during expiration resulted in a delay in the onset of the next inspiration, 3) During *in situ* baroreceptor stimulation, activity of the post-I neurons is increased, and aug-E activity is decreased in BötC, and 4) Finally, the model shows that baroreceptor input to post-I neurons of the respiratory CPG may be responsible for the effect while RSA has no influence on CVC. Taken together, the data support the hypothesis that the HB, by way of pulsatile baroreflex activation, controls the initiation of inspiration. This occurs through a rapid neural activation loop from the carotid baroreceptors to BötC expiratory neurons and the phrenic nerve in only a few synapses.

## Material and Methods

### Human Subjects

Subjects were young, healthy, yoga-naive males (N = 10, mean age 26.7 ± 1.4). A subset of data from this same subject pool was previously reported for analysis of blood pressure variability. Screening, consenting procedures, and details of instrumentation are the same as already reported (Dick *et al*., 2014b). Briefly, subjects were in the supine position during consecutive 20-min epochs of baseline breathing, uncoached slow deep breathing (SDB), and recovery breathing. Continuous monitoring was done for catheter-based brachial artery blood pressure, Lead II electrocardiogram, and calibrated double pneumobelt. The experiments and procedures were approved by the Institutional Review Board at the Mayo Clinic and conformed to the Declaration of Helsinki. All subjects signed an approved informed consent form. The data were de-identified to comply with HIPAA rules and regulations for data analysis. Further de-identification permitted data sharing without additional IRB approval.

### Rats (in situ preparation)

Rats (male, juvenile 50-100g) were pretreated with heparin sodium (1000 units, i.p.) and deeply anesthetized with isoflurane, bisected sub-diaphragmatically. We placed the rostral half of the rat in a cold (10°C) Ringer solution (containing, mm: NaCl, 125; NaHCO3, 25; KCl, 3; CaCl2, 2.5; MgSO4, 1.25; KH2PO4, 1.25; and dextrose, 10), where they were decerebrated pre-collicularly and had their skin, viscera, the left ribcage, diaphragm, lungs, and thoracic connective tissue removed; and then finally, the distal end of the descending aorta was freed for the perfusion cannula and the left phrenic was dissected free of connective tissue and desheathed for recording. The *in situ* preparation was moved to a recording chamber, cannulated and perfused retrogradely through the descending aorta with a modified Ringer’s solution (artificial cerebrospinal fluid - aCSF) saturated with 95% O_2_/5% CO_2_, and paralyzed with vecuronium bromide. Perfusion pressure was adjusted by manipulation of peristaltic pump’s rotation speed and by administration of supplemental vasopressin. After placement of the peripheral nerves in the recording electrodes, respiratory efforts were re-established by gradually increasing perfusion pressure and temperature. Motor activity patterns were recorded from the central end of the vagus, thoracic sympathetic and phrenic nerves.

The multi-electrode array was fitted to an electrode manipulator, which fit a stereotaxic frame. The microelectrodes (n=16, 10–12 MΩ) were aligned perpendicularly to the dorsal medullary surface. We placed eight electrodes bilaterally in two rows of four that paralleled the midline. The electrodes in the two rows were separated 250-μm, while electrodes within each row were separated by 300 μm. We used stereotaxic coordinates to position electrodes bilaterally in the rostral lateral medulla. We could position the depth of each electrode in steps as small as a micron and could adjust the electrode to optimize signal-to-noise ratio and to isolate the recording of activity to a single source. We characterized neuron recording by the peak of their activity during the respiratory cycle and the stereotaxic location of the electrode tip. In cases where more than one neuron was recorded on a single electrode, we discriminated single units using a voltage threshold and then confirmed single units using principal component analysis (spike sorting). The protocol included at least a 15-min baseline recording followed by characterizing the responses to transient increases in the perfusion pressure to activate arterial baroreceptors with 3-5 min interval between repeated activations of the baroreceptors.

We performed data analysis off-line from the Spike-2 files. Data were filtered from 100 to 3 kHz and the analog signal was sampled at 10jHz. The recorded data include: PNA, tSNA, ECG and extracellular potentials from the microelectrode array. PNA was processed by removing DC offset, rectification and smoothing using a 50-ms time constant to obtain a moving-time average of activity. From this ‘integrated’ PNA, we marked the onsets of inspiratory and expiratory phases. Action potentials of single neurons were converted to times of occurrence, *i*.*e*. spike trains (Fig. 5B).

### Analysis of human physiological data

The detection of respiratory phase changes in human ventilatory data in this dataset has been previously described (Barnett *et al*., 2020). Here, we constructed a probability density function (PDF) for heartbeats relative to the onset of inspiration for each of the three experimental epochs in each participant by collecting the times of each R-peak that happened within the 0.5 s interval immediately preceding the onset of inspiration. Each PDF was normalized and then integrated to produce a heartbeat cumulative distribution function (CDF), which could then be analyzed. We used the Kolmogorov-Smirnov test to determine whether CVC was present in these physiological recordings: the heartbeat cumulative distribution CDF was compared to the CDF of the uniform distribution.

In order to characterize the distribution of heartbeats preceding the onset of inspiration, we produced two metrics. For the first metric, we detected the maximum positive difference between the heartbeat CDF and the CDF of the uniform distribution. For the second metric, we recorded difference between the onset of inspiration and the time where the positive difference between the heartbeat CDF and the CDF of the uniform distribution was maximal. For the statistical analysis of these metrics, comparisons among groups were carried out using the CAR and PMCMRplus libraries for the R computing environment. Comparisons among the three experimental epochs were performed using a one-way repeated measures ANOVA or if group data were not normal with the Friedman test. In neither case (Fig. 3) were these comparisons significant, and post-hoc tests were not performed.

### Analysis of rat recordings

In rat recordings, we quantified the change in respiratory phase duration and the change in neuronal firing rate in cycles during which a baroreflex stimulation was performed. We designated cycles during which there was no stimulation as control cycles.

We analyzed and compared neuronal firing rates between control and perturbed cycles. For expiratory neurons in control cycles, we averaged the firing rate of neurons during expiration. Since the expiration duration was altered in perturbed cycles, we averaged the firing rate of neurons over the time interval that began at the beginning of expiration and ended after the average duration of expiration for control cycles in that cell. For inspiratory neurons in both control cycles and perturbed cycles, we averaged the firing rate of the cell over the duration of inspiration. We compared neuronal firing rates for control vs perturbed cycles using the paired t-test. The threshold for significance was 0.05.

We analyzed and compared respiratory phase durations between control and perturbed cycles. We separated the expiratory phase into the post-inspiratory phase and the late expiratory phase. The post-inspiratory phase lasted for 20% of the duration of the average expiration duration for control cycles. We compared respiratory phase durations between control and perturbed cycles using the Wilcoxon signed-rank test. The threshold for significance was 0.05.

Analysis and comparison of rat data were performed in Python using the Numpy, Scipy, and Pandas libraries.

### Model description

We developed two computational models of the brainstem respiratory circuitry: a simple model of baroreceptor stimulation in the rat, and a closed-loop model of blood-pressure derived baroreceptor activation in human beings. These two models shared fundamental core connectivity of the respiratory neuronal populations in the Bötzinger and pre-Bötzinger complexes, which was informed by brainstem transection experiments (Smith *et al*., 2007). As in (Rubin *et al*., 2009; Rubin *et al*., 2011), the model of the respiratory circuitry produced an average membrane potential for each neuronal population, which was transformed into the firing rate of that population.

### Model of rodent baroreceptor activation

By incorporating some critical slow intrinsic ionic conductances, this model captures the experimentally observed firing rate profiles of respiratory neurons. The average membrane potential of each neuronal population was determined by the following current balance equation:

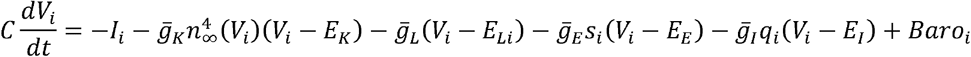

From *V*_*i*_, we computed the firing rate for each neuronal population with the piecewise-linear function *f*(*V*):

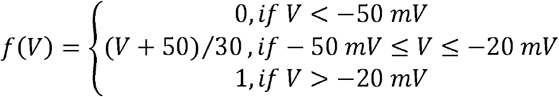

All five of the neuronal populations possessed a delayed rectifier potassium current and a leak current. The leak current was parameterized by its conductance, 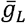, and its reversal potential, *E*_*L*_. The delayed rectifier potassium current was parameterized by its conductance, 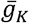, and the reversal potential of potassium, *E*_*K*_. The steady state activation of this potassium current was expressed as *n* _*∞*_(*V*) =1 / (1+exp (− (*V* + 30) / 4))

In the current balance equation, the additional intrinsic currents represented by *I*_*i*_ differed by neuronal population and defined the firing rate responses of each neuronal population to synaptic input. In the pre-I/I population (*I* = 1),*I*_*i*_ was composed of a slowly inactivating persistent sodium current 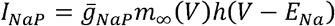. The current was parameterized by its maximal conductance, 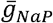, and the sodium reversal potential, *E*_*Na*_. The activation of its conductance was defined by its steady state voltage dependence:

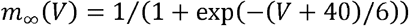

The inactivation of the persistent sodium current, *h*, was defined by the differential equation:

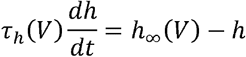

The steady state voltage dependence of *h* was defined by *h* _∞_ (*V*) = 1/ (1 + exp ((*V* + 55 / 10)), and its time constant was expressed as *τ* _*h*_ (*V*) = *τ*_*Nap*_ / cosh ((*V* + 55) /10). population (*i = 2*),*I* _*i*_ was composed of an adaptive potassium current 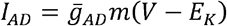, which was parameterized by its maximal conductance, 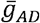, and the reversal potential of potassium, *E*_*K*_. The activation of *I*_*AD*_ was determined by the differential equation

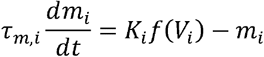

The steady state activation of *I*_*AD*_ was determined by the firing rate of the population and the scaling factor *K* _*i*_. In the aug-E population (*i* = 3), there were no additional intrinsic currents (*I*_*i*_ = 0). In the post-I population (*i* = 4) *I*_*i*_ was composed of an adaptive potassium current 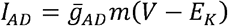; its dynamics are described above. This neuronal population also possessed the baroreceptor input stimulus, *Baro*, which was defined by the following differential equation:

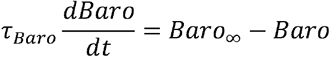

The parameter values for *Baro* _∞_ and τ_*Baro*_ were 0 nA and 1 s, respectively, while baroreceptor activation was absent. The values for biophysical parameters are given in Table 1. The firing rate of each population determined the instantaneous conductance of its synaptic current in post-synaptic populations. The excitatory (*S* _*i*_) and inhibitory (*q* _*i*_) synaptic activations were determined by the activity of presynaptic populations as described by

**Table 1.**
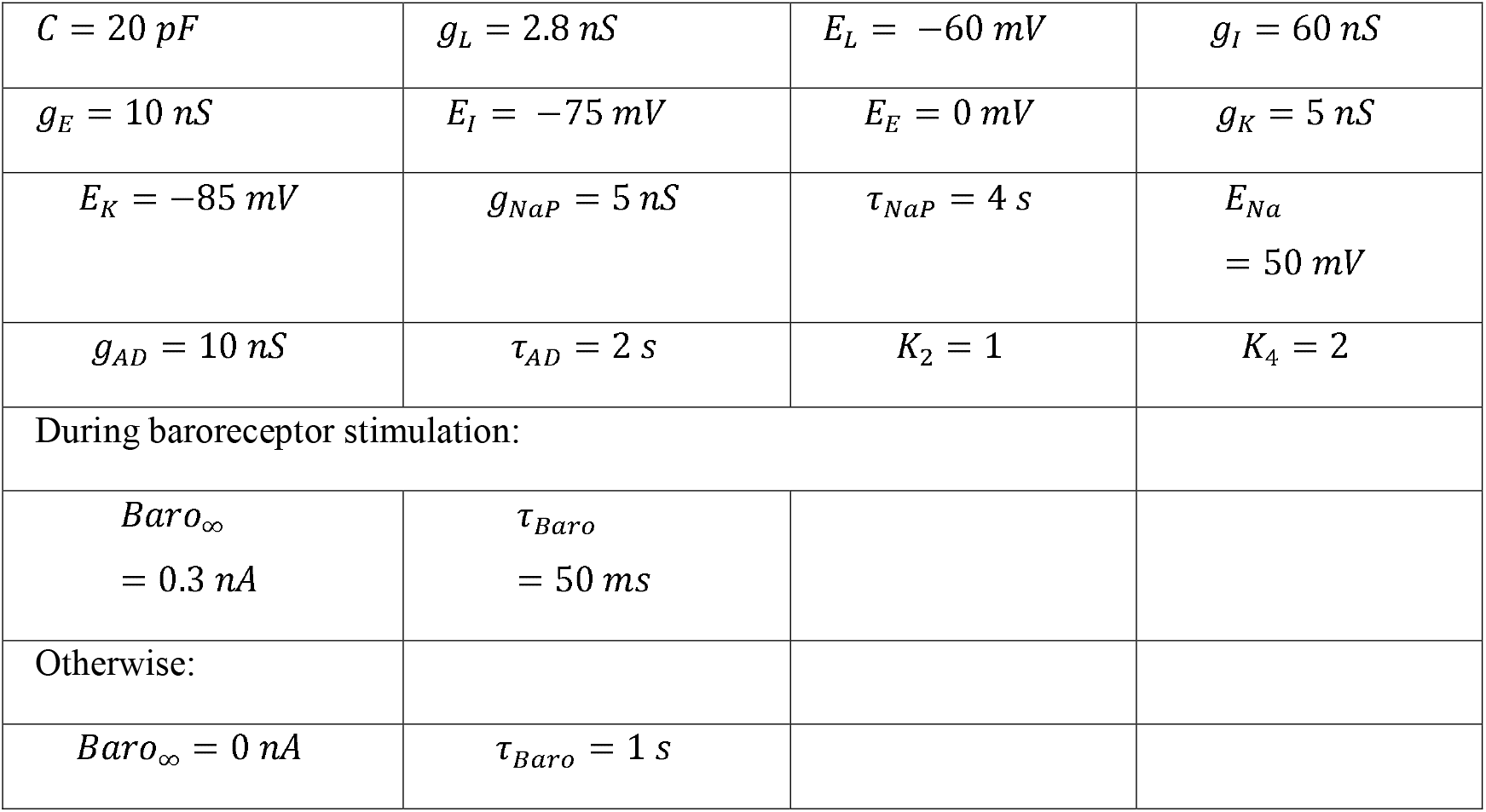
Values for parameters for the model of rodent baroreceptor activation.

|  |  |  |  |
| --- | --- | --- | --- |
| $C = 20 \text{ pF}$ | $g_L = 2.8 \text{ nS}$ | $E_L = -60 \text{ mV}$ | $g_I = 60 \text{ nS}$ |
| $g_E = 10 \text{ nS}$ | $E_I = -75 \text{ mV}$ | $E_E = 0 \text{ mV}$ | $g_K = 5 \text{ nS}$ |
| $E_K = -85 \text{ mV}$ | $g_{NaP} = 5 \text{ nS}$ | $\tau_{NaP} = 4 \text{ s}$ | $E_{Na} = 50 \text{ mV}$ |
| $g_{AD} = 10 \text{ nS}$ | $\tau_{AD} = 2 \text{ s}$ | $K_2 = 1$ | $K_4 = 2$ |
| During baroreceptor stimulation: |  |  |  |
| $Baro_{\infty} = 0.3 \text{ nA}$ | $\tau_{Baro} = 50 \text{ ms}$ | | |
| Otherwise: |  |  |  |
| $Baro_{\infty} = 0 \text{ nA}$ | $\tau_{Baro} = 1 \text{ s}$ | | |

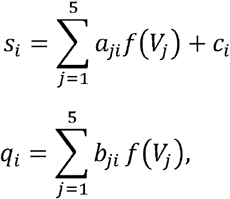

where *f* (*V*_*j*_) is the neuronal firing rate of the presynaptic population. The synaptic weight *a*_*ji*_ corresponds to the specific strength of the excitatory projection from population *j* to population *i*. The synaptic weight *b*_*ji*_ corresponds to the specific strength of the inhibitory projection from population *j* to population *i*. The synaptic weight *c*_*i*_ represents a tonic excitatory drive to population *i*. The magnitude of these weights can be found in Table 2. Simulations were performed in MATLAB using the ode15s solver (AbsTol = 1e-7, RelTol = 1e-5, and MaxStep = 10).

**Table 2.** Weights of synaptic connections among neuronal populations.

|  | Pre-I/I | Early-I | Aug-E | Post-I | Drive |
| --- | --- | --- | --- | --- | --- |
| Synaptic | $a_{1i}$ | $b_{2i}$ | $b_{3i}$ | $b_{4i}$ | $c_i$ |

| Weights |  |  |  |  |  |
| --- | --- | --- | --- | --- | --- |
| Pre-I/I |  |  | 0.15 | 1 | 0.02 |
| Early-I | 0.15 |  | 0.05 | 0.42 | 0.6 |
| Aug-E |  | 0.42 |  | 0.25 | 0.45 |
| Post-I |  | 0.22 |  |  | 0.6 |

### Model of human cardiorespiratory interaction

We extended and adapted our model of rodent baroreceptor activation to simulate human cardiorespiratory interaction by incorporating the HB, blood pressure, and the tidal volume of the lungs.

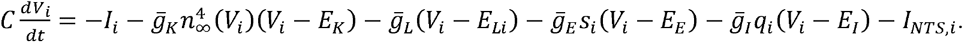

This model included an additional neuronal population, which was responsible for the inspiratory output of the respiratory CPG. The ramp-I population (*i* = 5) possessed a leak current, and a delayed rectifier potassium current, but there were no additional intrinsic currents (*I*_*i*_ = 0). The intrinsic currents of the pre-I/I, early-I, and aug-e populations are identical to those described inthe model above. The post-I population (*i* = 4) differed from the previous model. It possessed the outward adaptive potassium current (*I*_*AD*_) as well as input from the Nucleus of the Solitary Tract (NTS).

We defined the dynamic input to the respiratory CPG from the NTS as *I*_*NTS,i*_. The only population receiving this input was the post-I populations (*i* = 4). This input was composed of synaptic currents related to the Hering-Breuer reflex, *I*_*HB*_, and the carotid baroreflex, *I*_*CB*_, such that *I* _*NTS*_ = *I*_*HB*_ +*I*_*CB*_ The Hering-Breuer reflex was defined as *I*_*HB*_ = *α*_*HB*_ *L* where _B_ was a gain parameter and was the tidal volume (is defined below). The carotid baroreflex was defined as

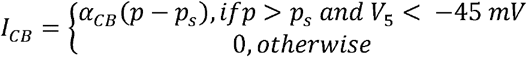

Where *α*_*CB*_ was a gain paremeter,*p* is the blood pressure (defined below), is the smoothed blood pressure (defined below), and *V*_5_ is the average membrane potential of the ramp-I population.

The tidal volume of the lungs is defined by

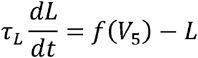

such that *f*(*V*_5_) is the firing rate of the ramp-I neuronal population. The blood pressure is defined by the differential equation

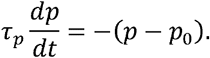

The steady state blood pressure was *p*_0_.=(*p*_1_− *p*_2_ *exp* (−1/(Ω *τ*_*p*_))) / (1−*exp* (−1 / (Ω *τ*_*p*_))), where *p*_1_ and *p*_2_ are nominal systolic and diastolic pressures, Ω is nominal heart rate, and *τ*_*p*_ is blood pressure relaxation constant. We also utilize a smoothed blood pressure approximately representing average blood pressure over time *τ*_*s*_. It was defined by the differential equation

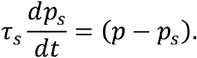

The activation of the adaptive potassium current (*I*_*AD*_) was altered in this model to account for variability in the duration of inspiration and expiration. The activation variable was defined as a stochastic differential equation:

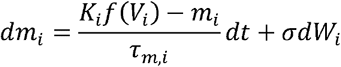

Where *W*_*i*_ is Wiener stochastic process, and the magnitude of *σ* defined respiratory variability.

The beating of the heart was described by non-homogenous Poisson process with the rate

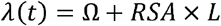

Where *L* is the tidal volume, and *RSA* is a parameter that defines the amplitude of respiratory sinus arrythmia. To approximate the heart rate variability observed in human data, the cardiac cycle was divided into 200 states, and the transition time between states was generated as an exponentially distributed random number with rate *λ*(*t*)×200.

Biophysical parameters and synaptic weights specific to the model of human cardiorespiratory interaction can be found in Tables 3 and 4. This model was used for simulations of regular restful breathing as well as slow deep breathing. In order to increase the duration of inspiration and expiration to mimic slow deep breathing in humans, we decreased the tonic excitatory drive to the early-I (*i* = 2) and post-I (*i* = 4) neuronal populations as well as the amplitude of the Weiner process (values given in Tables 3 and 4). Simulations were performed using custom software written in C++. Differential equations were solved using the stochastic Euler-Maruyama method with a step of 0.1 ms.

**Table 3.**
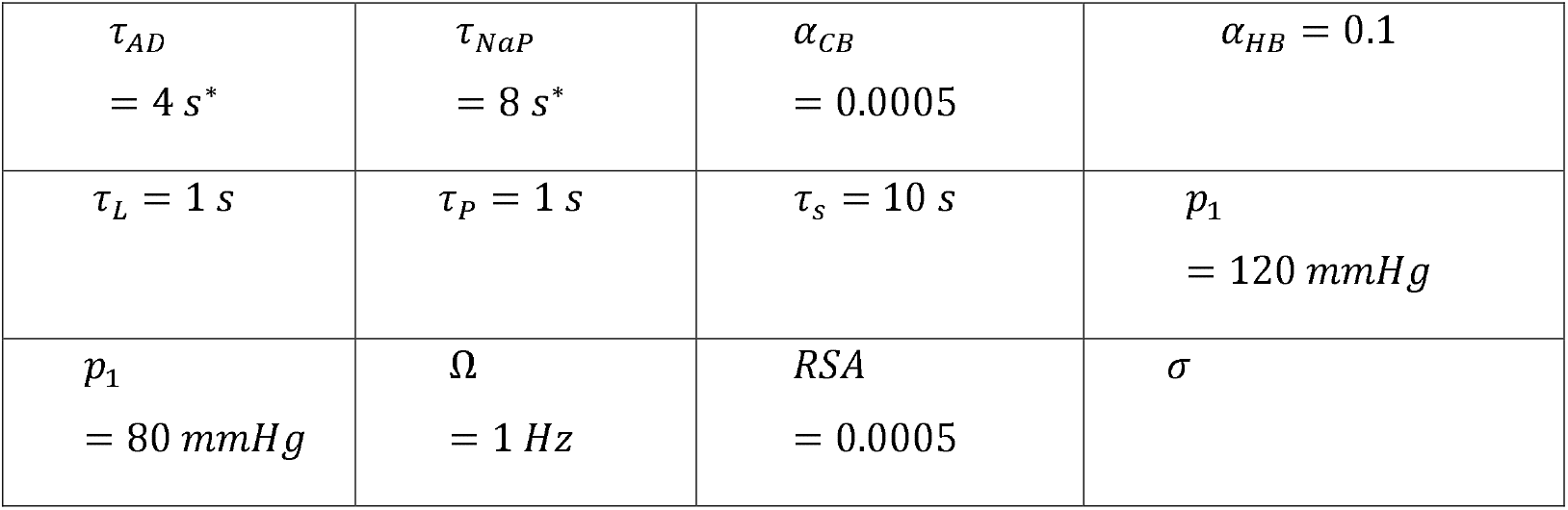
Summary of biophysical parameters for the model of human cardiorespiratory interaction. Quantities marked with an asterisk emphasize values that differ from the model of rodent baroreflex stimulation. Quantities contained within parentheses indicate the parameter value used for simulations of slow deep breathing.

|  |  |  |  |
| --- | --- | --- | --- |
| $\tau_{AD}$<br>= 4 s* | $\tau_{NaP}$<br>= 8 s* | $\alpha_{CB}$<br>= 0.0005 | $\alpha_{HB} = 0.1$ |
| $\tau_L = 1$ s | $\tau_P = 1$ s | $\tau_s = 10$ s | $p_1$<br>= 120 mmHg |
| $p_1$<br>= 80 mmHg | $\Omega$<br>= 1 Hz | $RSA$<br>= 0.0005 | $\sigma$ |

**Table 4.**
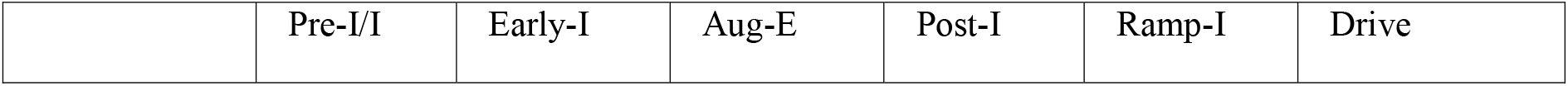
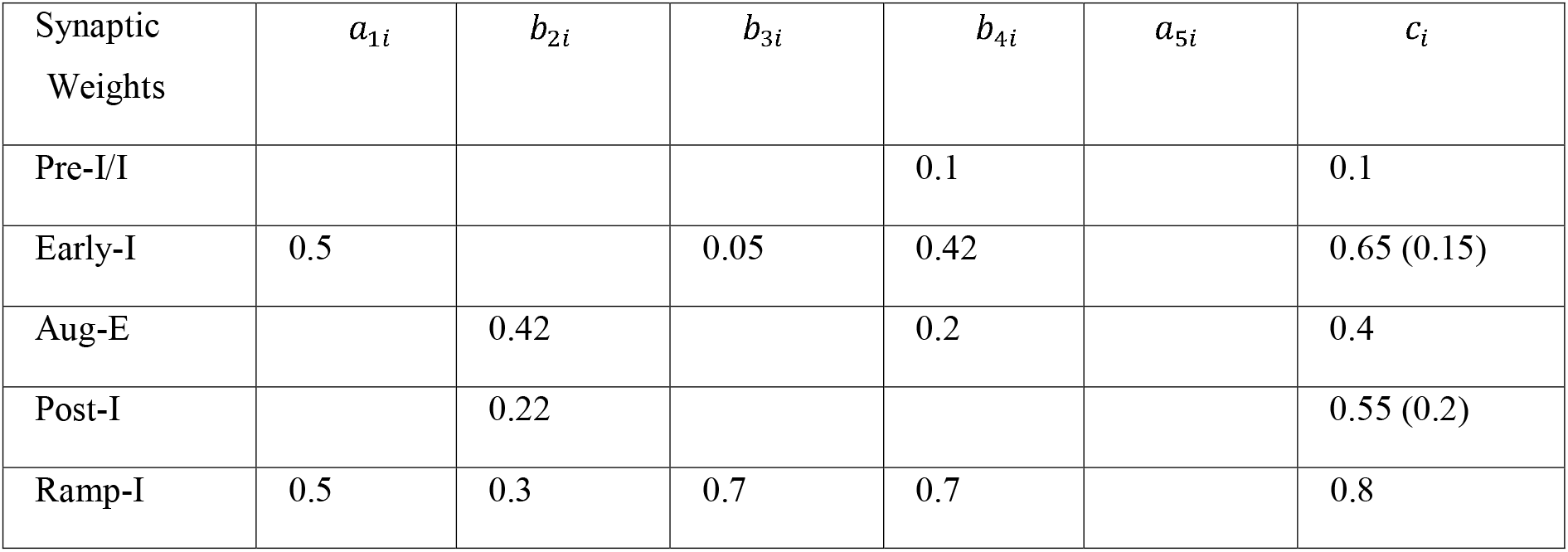
Summary of synaptic connectivity among neuronal populations for the model of human cardiorespiratory interaction. Quantities contained within parentheses indicate the parameter value used for simulations of slow deep breathing.

| Synaptic Weights | $a_{1i}$ | $b_{2i}$ | $b_{3i}$ | $b_{4i}$ | $a_{5i}$ | $c_i$ |
| --- | --- | --- | --- | --- | --- | --- |
| Pre-I/I |  |  |  | 0.1 |  | 0.1 |
| Early-I | 0.5 |  | 0.05 | 0.42 |  | 0.65 (0.15) |
| Aug-E |  | 0.42 |  | 0.2 |  | 0.4 |
| Post-I |  | 0.22 |  |  |  | 0.55 (0.2) |
| Ramp-I | 0.5 | 0.3 | 0.7 | 0.7 |  | 0.8 |

## Acknowledgments

The study was supported by NIH grants R01 AT008632 (Y.M.), U01 EB021960 (T.D. and Y.M.). The human data was collected by E.W. in collaboration with Michael Joyner under support of the NIH grant HL 083947. JFRP is supported by the Marsden Fund Council from Government funding, managed by Royal Society Te Apārangi, and Health Research Council of New Zealand.

